# Distinct Functional Roles of Narrow and Broadband High-Gamma Activities in Human Primary Somatosensory Cortex

**DOI:** 10.1101/2024.03.18.585646

**Authors:** Seokyun Ryun, Chun Kee Chung

## Abstract

In previous studies, higher (broadband) and lower (narrowband) components of high-gamma (HG) activity (approximately from 50 to 150 Hz) have different functions and origins in the primary visual cortex (V1). However, in the primary somatosensory cortex (S1), it is unknown whether those are similarly segregated. Furthermore, the origin and functional role of S1 HG activity still remain unclear. Here, we investigate their roles by measuring neural activity during vibrotactile and texture stimuli in humans. Also, to estimate their origins, S1 layer-specific HG activity was measured in rats during somatosensory stimulation. In the human experiment, with texture stimulation, the lower HG activity (LHG, 50-70 Hz) in S1 represents the intensity of the sustained mechanical stimulus. In the vibrotactile experiment, the higher HG (HHG, 70 -150 Hz) activity in S1 depended on the ratio of low and high mechanical frequencies with its pattern being a mixture of neural activity for low and high mechanical frequencies. Furthermore, 8 texture types could be classified using power values of HHG activity, while the classification using LHG activity showed poor performance. In the rat experiment, we found that both HHG and LHG activities are highest in the somatosensory input layer (layer IV), similar to previous visual cortex studies. Interestingly, analysis of spike-triggered LFP (stLFP) revealed significant HG oscillations during pressure stimulation with the stLFP HG power most significant in layer IV, suggesting that both LHG and HHG activities are closely related to the neuronal firing in layer IV. In summary, LHG activity represents the intensity of tactile sensation, while HHG activity represents the detail of the surface geometry of objects interacting with skin. Additionally, low and high mechanical frequencies are processed in parallel in S1. Finally, both HHG and LHG originated in layer IV of S1.

## INTRODUCTION

High-gamma (HG) band (approximately from 50 to 150 Hz) activity is one of the most robust neural signals recorded in invasive electrophysiological recording, including electrocorticography (ECoG). In the human somatosensory system, the HG activity has been consistently observed during somatosensory stimulation not only in the primary somatosensory cortex (S1) but also in the downstream areas, including secondary somatosensory cortex (S2), posterior parietal cortex (PPC) and premotor cortex (PM) (Avanzini et al., 2016; Ryun et al., 2017b; Ryun et al., 2023). In our previous study, we suggested that this activity in somatosensory-related areas may be related to the perceptual processing of somatosensation (Ryun et al., 2023). However, its mechanisms and functional roles have been poorly understood even in S1. Specifically, its relationship to conventional gamma activity also remains unclear (Crone et al., 2011).

In vision studies, it has been suggested that broadband (∼80 to 200 Hz) and narrowband gamma (close to 60 Hz) activities in the primary visual cortex (V1) have different origins and functions in mice and monkeys (Ray and Maunsell, 2011; Saleem et al., 2017). In particular, a recent study indicated that narrowband gamma activities (50 to 70 Hz) are closely associated with neuronal firing in the lateral geniculate nucleus (LGN) of the thalamus and propagate throughout the thalamocortical visual system, including V1 and higher visual areas (Shin et al., 2023). A human vision study using ECoG suggested that narrowband gamma (∼20 to 60) and broadband gamma (∼70 to 150+ Hz) activities showed distinct spectral, temporal, and functional properties (Bartoli et al., 2019). However, although some somatosensory studies indicated that HG activity represents specific physical properties of delivered stimuli, its exact functional roles still remain to be elucidated. Furthermore, it is unclear whether broadband HG and narrowband gamma activities in the S1 can be functionally dissociated as in the visual cortex (Jiang et al., 2020; Ryun et al., 2017b).

The generation mechanisms of conventional gamma oscillatory activity are relatively well established especially in the hippocampus and thalamocortical network (Fernandez-Ruiz et al., 2023; Ray and Maunsell, 2015; Saleem et al., 2017; Whittington et al., 1995). For example, in hippocampus CA1, there exist three different gamma generation mechanisms (30-50 Hz, 60 to 100 Hz, and 100 to 150 Hz gamma oscillations) which can be segregated by their micro-level circuit structures (Buzsáki and Wang, 2012; Fernandez-Ruiz et al., 2023). Additionally, in the sensory cortex, the unidirectional flow of neural signals from the thalamus to the cortex is closely related to the narrowband gamma oscillation (Saleem et al., 2017). In contrast, although there is a finding that broadband HG activity is tightly linked to neuronal spikes (Ray et al., 2008a), the mechanism of HG activity and why this activity is tightly associated with neuronal firing have been poorly understood.

The exact frequency boundaries of gamma and HG activities are ambiguous. The lower boundary of HG activity has often been considered to be 50 to 80 Hz (Avanzini et al., 2018; Canolty et al., 2006; Ray et al., 2008b; Ryun et al., 2017a), but this appears to vary between individuals and across cortical regions. Likewise, it is difficult to clearly define the upper boundary of gamma and broadband HG activities. Given that these boundaries show high individual and regional variability, strict frequency-based definitions of neural activity may be inadequate, at least in the gamma and HG ranges. Rather, a mechanism- or function-based definition of these activities may be more appropriate (Fernandez-Ruiz et al., 2023). However, paradoxically, to investigate their functional roles and mechanisms, it is inevitable to select the approximate frequency ranges of these activities. In this study, we set the HG frequency range in S1 to be 50 to 150 Hz for consistency with our previous studies.

In this study, we test the hypothesis that HG activities in S1 can be segregated by their functions in humans. Specifically, we focus on the lower (50 to 70 Hz) and higher (70 to 150 Hz) frequency components of HG band activity.

Furthermore, we investigate the origin of two HG activities by analyzing cortical layer-specific local field potentials (LFP) and neuronal firing in rat S1.

## RESULTS

Eight patients with subdural electrodes on the S1 hand area were included in this study. Among them, seven patients performed the vibrotactile stimulation task, and three patients participated in the texture stimulation task. The demographics of the patients are presented in Table 1.

In our previous studies, we consistently observed high-frequency neural activity approximately from 50 Hz in S1, while no significant activity was observed at the low gamma range (30 to 50 Hz) (Ryun et al., 2017a; Ryun et al., 2017b; Ryun et al., 2023). For consistency of our studies, in this study, we chose the HG frequency range as 50 to 150 Hz and divided it into two components: lower component of HG activity (LHG; 50 to 70 Hz) which is similar to the narrowband gamma oscillation in other studies, and higher component of HG activity (HHG; 70 to 150 Hz). The boundary of two components (70 Hz) was determined based on the frequency range of conventional gamma oscillation (30 to 70 Hz), a recent study (narrowband gamma oscillation: 50 to 70 Hz) (Shin et al., 2023), and visual inspection of our data. Note that slight changes in boundary frequency (65 to 75 Hz) did not significantly affect the main results.

### The lower component of HG activity represents the strength of sustained mechanical stimulus

To test whether HG activity represents specific functions depending on the frequency, we designed a texture stimulation paradigm with two different contact forces (light vs. heavy conditions) and eight different surface textures. We observed strong HG activities both in heavy and light conditions, but their spectro-temporal pattern showed distinct differences. Overall, we found three types of HG patterns during heavy/light texture stimulation. (1) a transient activity from 0 to 250 ms in the frequency range of 70-150 Hz (transient HHG, tHHG); (2) an intermittent activity approximately from 250 ms to 1.5 s in the frequency range of 70-150 Hz (HHG); (3) a strong and sustained narrowband HG activity from 250 ms to 1.5 s (50 to 70 Hz; LHG) (**Figure 1**, top). Additionally, the LHG band showed long-latency tonic activity, while 70-150 Hz HG activity strikingly increased with very short latency and then decreased over time but still maintained significant activity compared to the baseline one (**Figure 1**, bottom). Interestingly, HHG activity appeared to be independent of the contact force delivered, even though both tHHG and LHG activities increased with increasing contact force. Specifically, LHG activity maintained a high activation level during the whole stimulus period, while tHHG was only observed at the on/offset of stimuli. Throughout the experiment, LHG activities of all three subjects significantly increased with increasing contact force of texture stimulation (**Figure 2**). In contrast, HHG activities during stimulation showed inconsistent patterns. Therefore, LHG activity in S1 is likely to be closely related to the intensity of sustained tactile stimulation.

**Figure 1.**
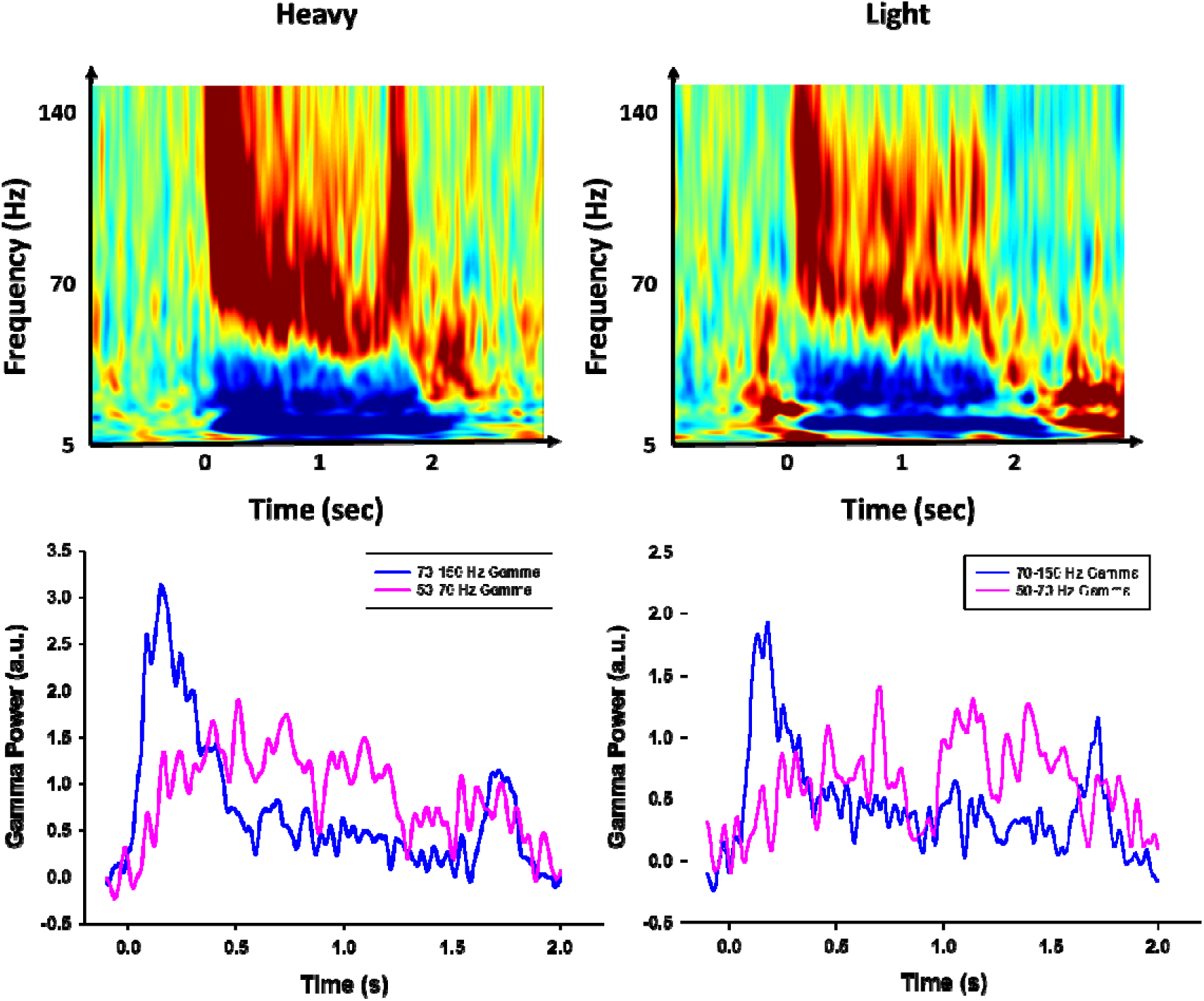
Representative time-frequency plots for texture stimuli under the heavy (top left) and light (top right) conditions and corresponding high-gamma time series plots (bottom). The bursting component (70-150 Hz during 0 to 250 ms period) and the low component (50-70 during 250 ms to 1.5 s) of high-gamma power levels increased with increasing normal force.

**Figure 2.**
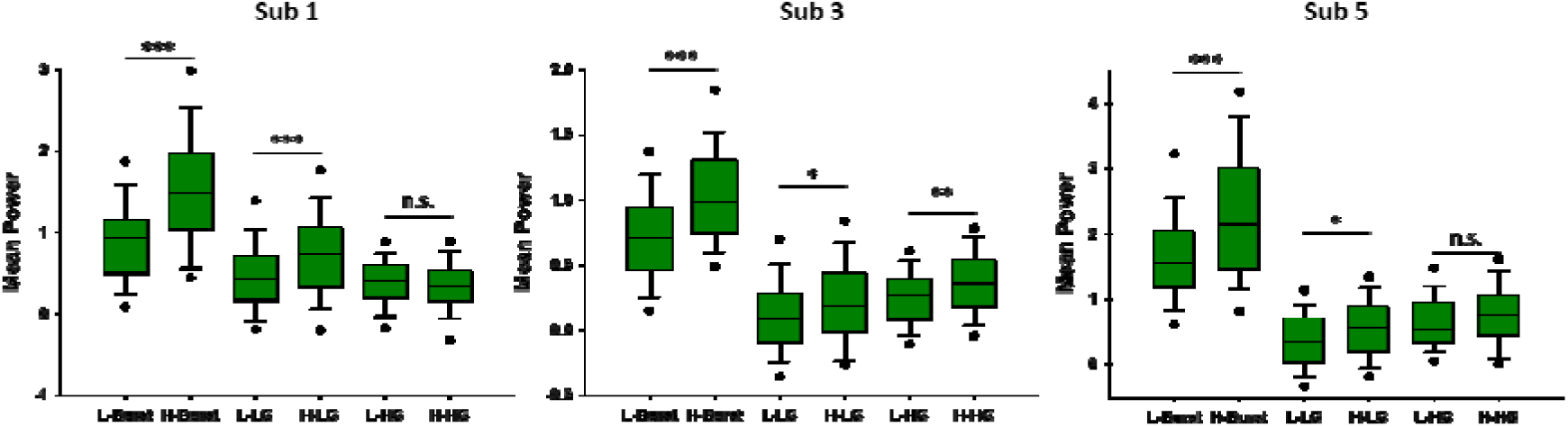
Mean power of each component under light and heavy conditions. L-burst indicates the mean power of transient HG activities (70-150 Hz during 0 to 250 ms) under the light condition, and H-burst indicates the power under the heavy condition. L-LG indicates the mean power of the LHG (50-70 Hz during 250 ms to 1.5 s) under the light condition (H-LG: heavy condition). L-HG indicates the mean power of the HHG activity (70-150 Hz) under the light condition (H-HG: heavy condition). The power levels of transient and LHG activities consistently and significantly increased with increasing normal force. In contrast, sustained HHG (70-150 Hz during 250 ms to 1.5 s) activities showed inconsistent patterns.

### HG activities for complex vibrotactile stimuli

Next, we focused on the functional role of HHG activity. In our previous study, we showed that the temporal pattern of HG activity varies depending on the vibrotactile frequency (Ryun et al., 2017b). That is, the rate of decline in HG activity increased by vibrotactile stimuli was faster in the high-frequency vibrotactile condition (>100 Hz) compared to the low-frequency condition (<100 Hz). However, it is unclear whether this phenomenon can be preserved when complex vibrotactile stimuli are delivered. To assess this issue, we delivered vibrotactile stimuli having various flutter (32 Hz) vs. vibration (350 Hz) magnitude ratios (99 (flutter):1 (vibration), 75:25, 50:50, 25:75, 1:99) to the patient’s contralateral index fingertip. We found that the pattern of HHG activity in S1 during complex vibrotactile stimulation was dependent on the magnitude ratio of flutter to vibration (**Figure 3 and Figure S1**). That is, the HHG pattern in the 75:25 condition was more similar to that in the 99:1 condition compared to that in the 1:99 condition. Conversely, the HHG pattern in the 25:75 condition was more similar to that in the 1:99 condition compared to that in the 99:1 condition. The sum of HHG power differences over time was the largest under the 99:1 vs. 1:99 condition, and these HHG pattern differences between the 99:1 condition and each condition showed a monotonically increasing trend with increasing vibration ratio and vice versa (**Figure 4A**). Furthermore, we found that the HHG activity of complex vibrotactile stimulation can be expressed as a superposition of HHG activities for flutter and vibration-only stimuli (**Figure 4B**). In other words, the HHG pattern of complex vibrotactile stimulation appeared to be a mixture of neural activities for flutter and vibration.

**Figure 3.**
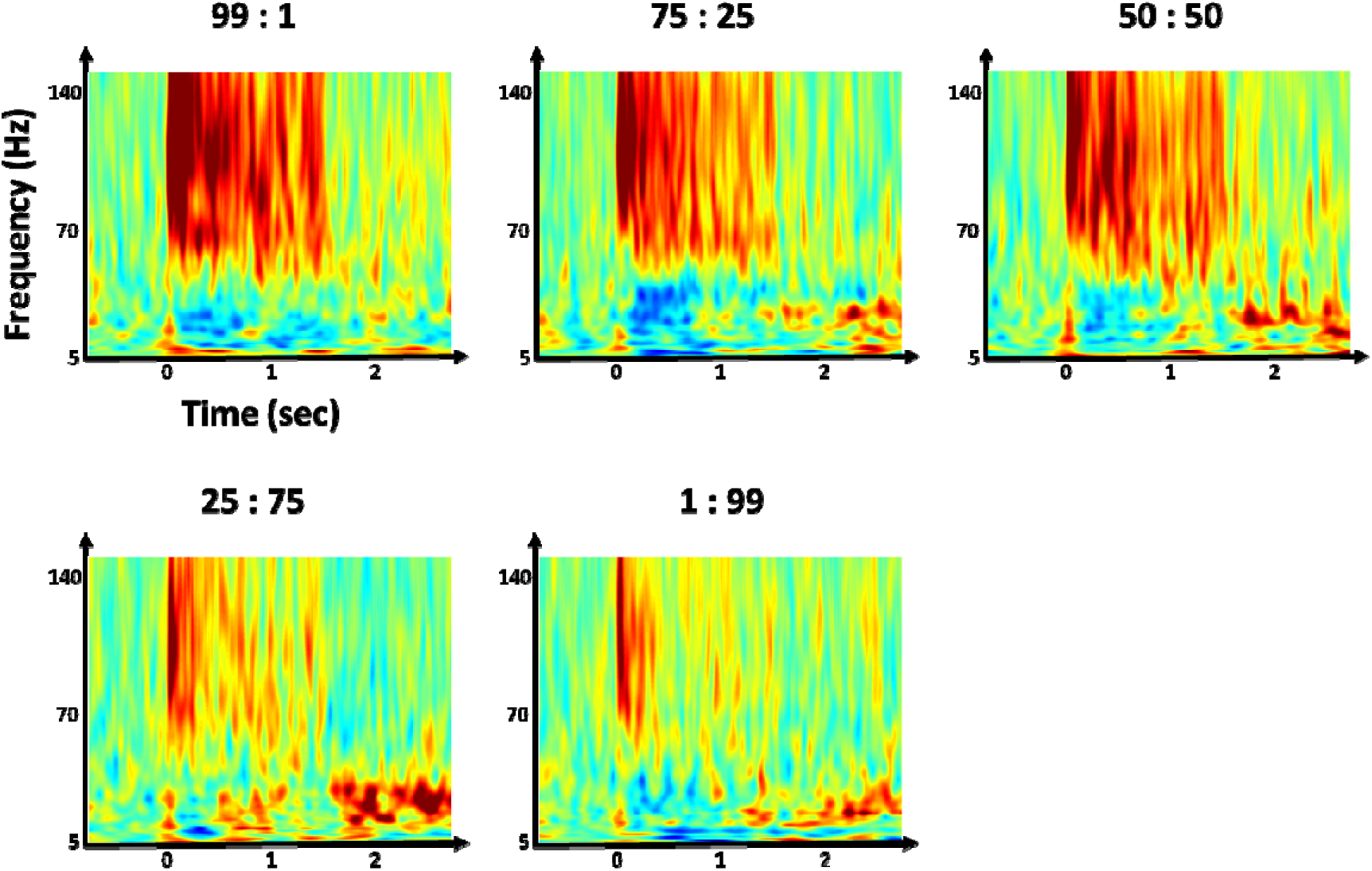
Time-frequency maps under various flutter (32 Hz) + vibration (350 Hz) conditions from subject 8. Legends above the maps indicate the ratio of flutter and vibration (e.g. 75:25 = flutter 75% and vibration 25%) based on the magnitude of vibrotactile stimulus. The maximum flutter magnitude (100%) was 120 μm and the maximum vibration one was 60 μm. Results show that sustained high-gamma activity becomes shorter when stimulation contains more vibration frequency (from 99:1 (flutter:vibration) to 1:99).

**Figure 4.**
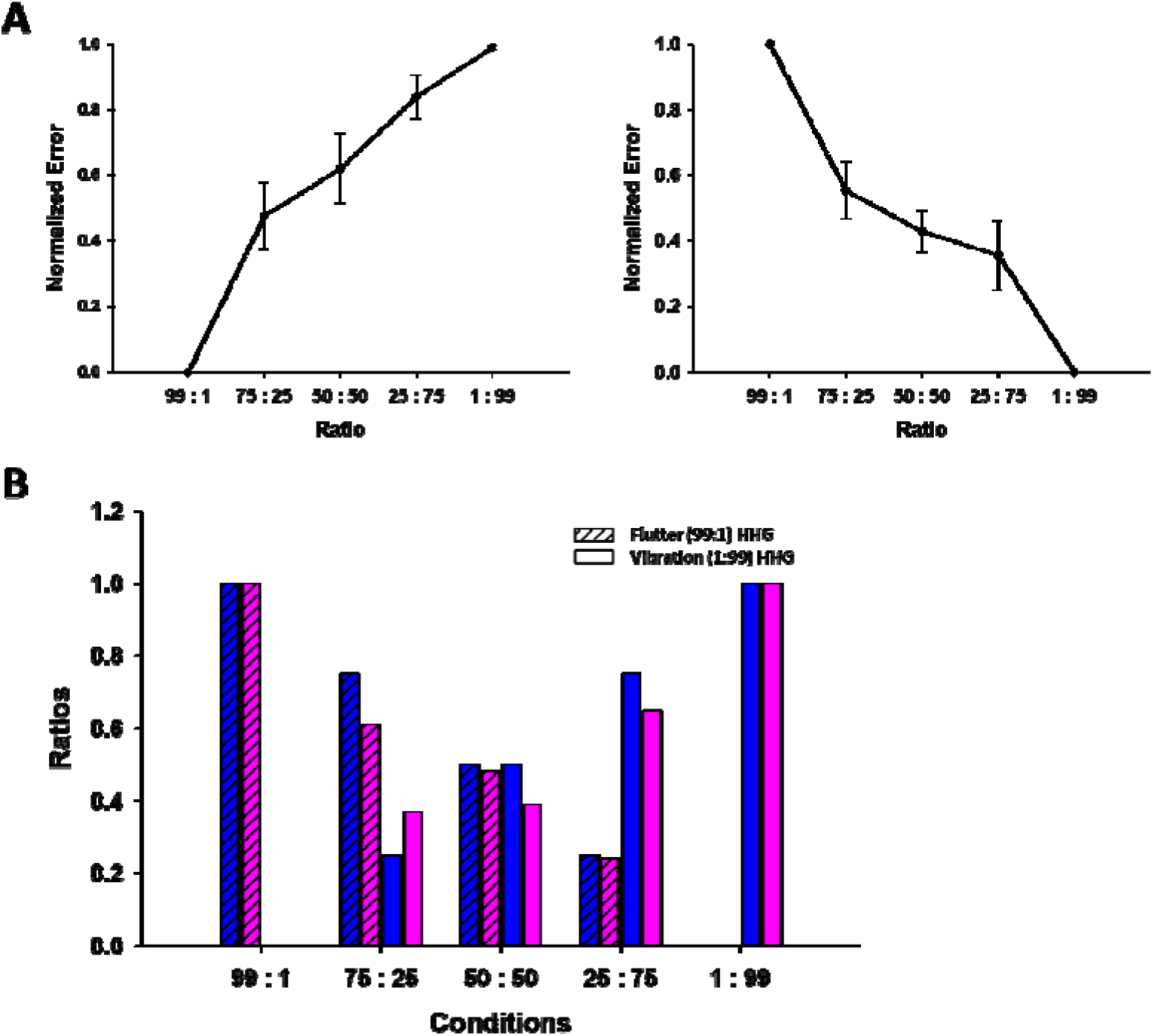
HHG pattern of complex vibrotactile stimulation is a mixture of neural activities for flutter and vibration. (A) High-gamma similarities (normalized error = 0 indicates that two stimuli are the same) between 1:99 vs. each stimulus (left) or 99:1 vs. each stimulus (right) condition during the stimulation period. Figures show monotonic error trends. Error values were obtained by calculating the difference in power at all time points during stimulation. The error values were then normalized based on the value of the maximum error condition. (B) HHG activity of complex vibrotactile stimulation can be expressed as a superposition of HHG activities for flutter and vibration-only stimuli. Blue bars indicate the reference values of each condition (e.g., for 75:25 condition). Pink bars indicate the estimated values (diagonal pattern: flutter-only HHG activity; blank: vibration-only HHG activity) of each condition (coefficients *a* and *b*, see Materials and Methods). For example, HHG activity for 25:75 complex vibrotactile stimulation can be expressed as a superposition of

### HHG activity represents the details of tactile information

Given the result from complex vibrotactile stimulation, we hypothesized that HHG activity during tactile stimulation may be related to the surface information of objects interacting with the skin. To test this, we delivered eight different textures to the patient’s fingertip and extracted features from the power time series of each HG component (tHHG, HHG, and LHG). The features of each component consisted of the power levels of five time bins (50 ms bins for transient HG (0 to 250 ms), and 250 ms bins for HHG and LHG (250 ms to 1.5 s)). We then tested whether the eight textures can be classified by these features. For classification, we used the customized multi-layer perceptron (MLP) classifier. Among the three conditions (tHHG, HHG, and LHG), the HHG condition was found to have the highest classification accuracy regardless of the contact force (light and heavy conditions; **Figure 5**). The highest classification accuracy for the HHG condition was 28.9 % for Subject 5 (chance level = 12.5 %). In contrast, the classification accuracy for the LHG condition was close to the chance level (overall accuracy = 12.14 %). These results indicate that HHG activity contains meaningful information about texture characteristics.

**Figure 5.**
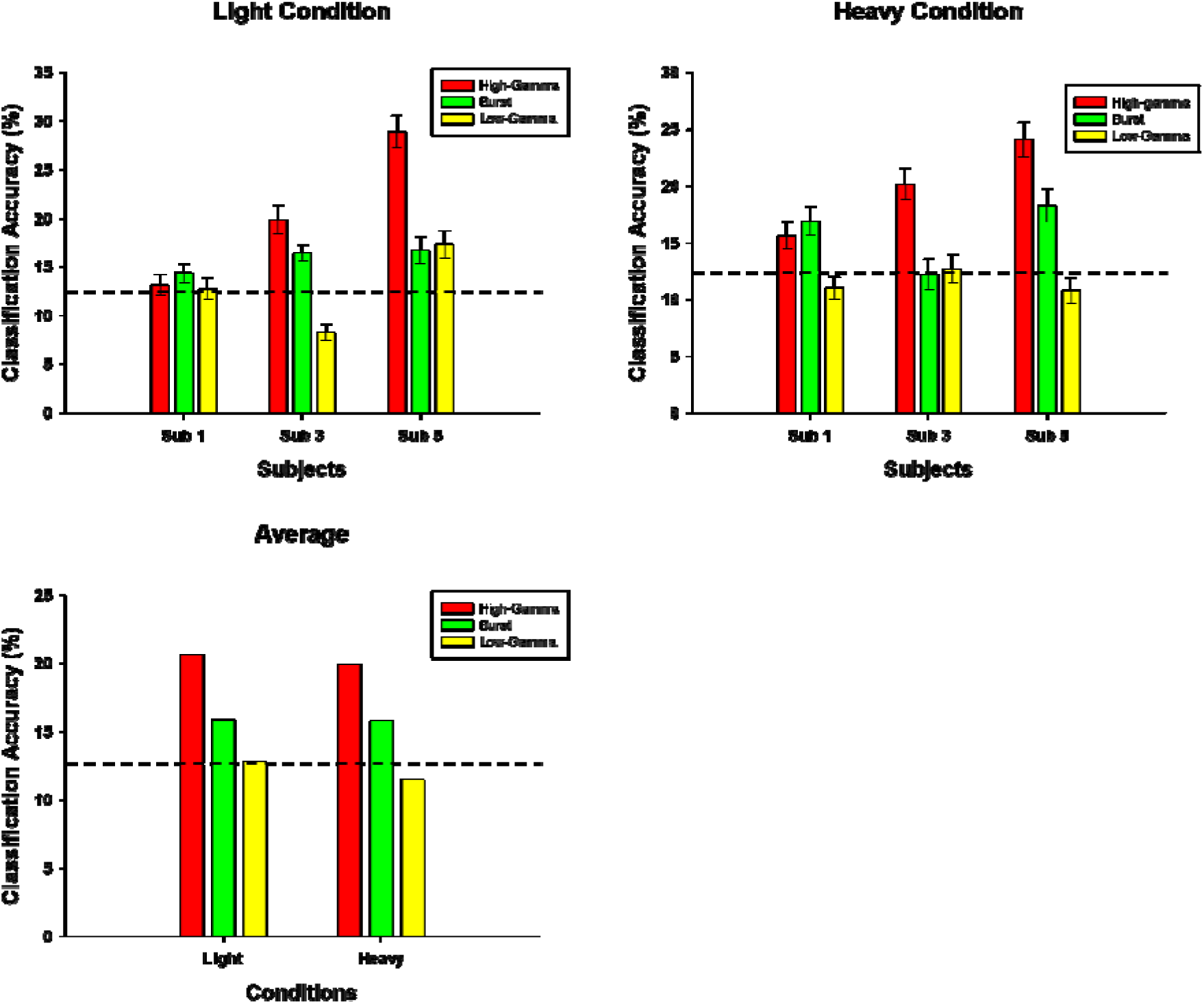
Classification accuracies for eight texture materials under light (top left) and heavy (top right) conditions. The bottom left figure indicates the average accuracy across subjects. Red bars indicate classification accuracy under the high component of HG activity condition (70-150 Hz during 250 ms to 1.5 s). Green bars indicate classification accuracy under the bursting activity of high-gamma (70 -150 Hz during 0 to 250 ms) condition, and yellow bars represent that under the low-component of high-gamma activity (50-70 Hz during 250 ms to 1.5 s) condition. Dashed horizontal lines indicate chance level (12.5 %). Error bars indicate the standard error of the mean (s.e.m.).

### Origin of HHG and LHG activities: The rat microelectrode study

Our results indicated that HHG and LHG activities are functionally different. However, it is unclear where these activities originate among cortical layers and whether they come from the same neural source. To assess this, we implanted a microelectrode bundle, which can record layer-specific neuronal activity, into the left S1 hindlimb region of a rat. We delivered ramp-and-hold pressure stimuli to the right hindlimb of the anesthetized rat (**Figure 6A** and **B**). Prior to analyzing layer-specific HG activity, we first confirmed the similarity of neural activities between rat and human during pressure stimulation (**Figure 6C** and **D**). We observed robust transient HG activities at the on/offset and sustained activities during stimuli in both rat and human. Through layer-specific HG analysis, we found that the power levels of both HHG and LHG activities are highest in layer IV, known as the major somatosensory input layer (**Figure 7A**). Next, we investigated whether these activities are related to neuronal spiking activity at each layer. To do this, we calculated the spike-triggered LFPs (stLFPs) of each electrode pair during stimulation. We observed significant HG band oscillatory activities in some spike-LFP pairs (electrode pairs 1 and 6; Figure 7C). Interestingly, both LHG and HHG band activities of stLFPs were most significant in layer IV (Figure 6D), suggesting that neuronal activity in the input layer is closely related to both LHG and HHG activities. However, we could not record the neural signal from the upper region of the superficial layer due to the insertion depth of the microelectrode bundle. Although it is possible that the upper region generates stronger HHG or LHG activity than layer IV, the current result indicates that layer IV is the major source of HHG and LHG activities among layers III-VI. At this point, further investigation is required to confirm our result.

**Figure 6.**
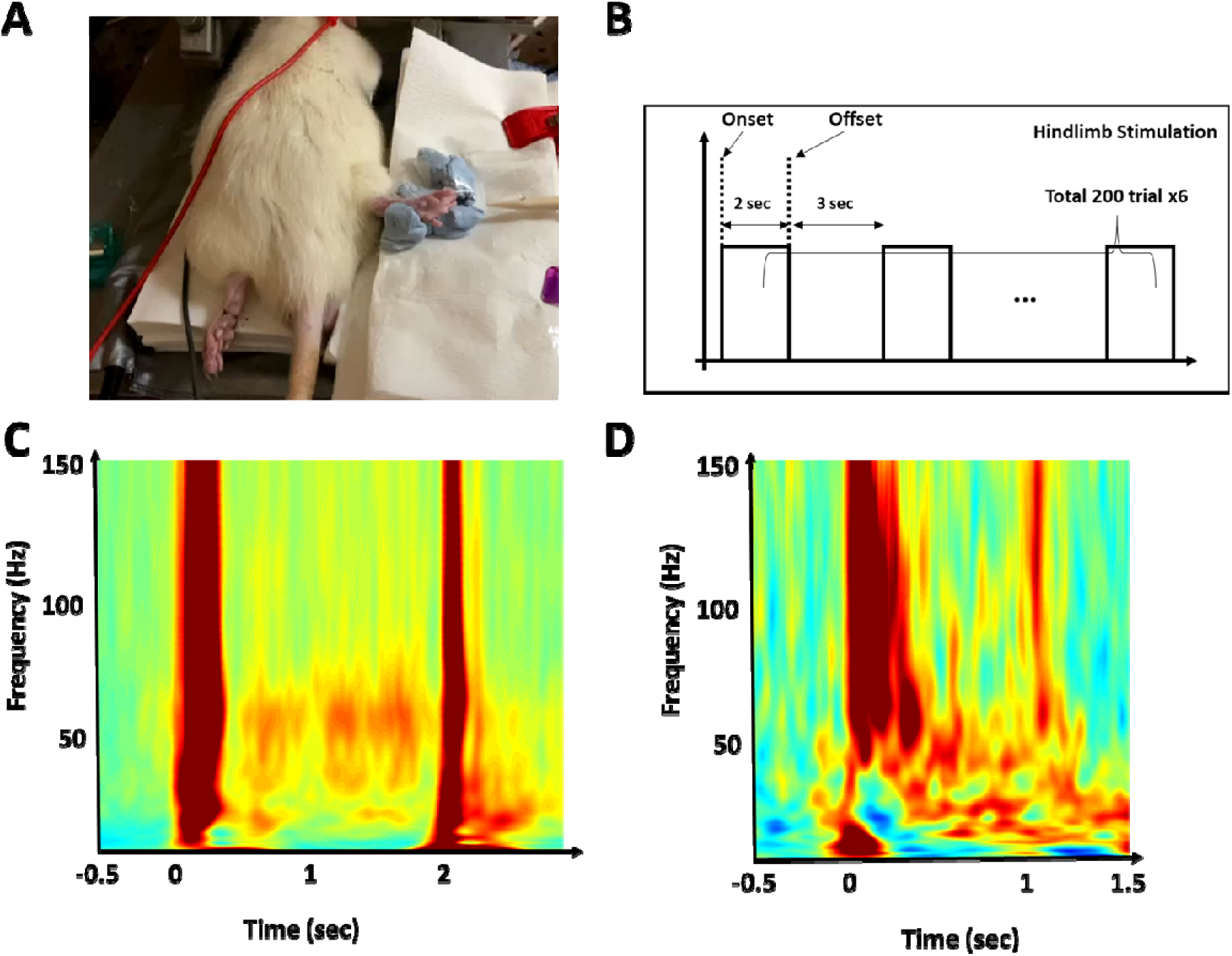
Rat experiment. (A) Pressure stimulation on anesthetized rat’s hind limb. (B) Stimulation paradigm. (C) Representative time-frequency plot for pressure stimulation in rat’s S1. (D) Time-frequency plot for pressure stimulation in human’s S1. Stimulus duration was approximately 1 s.

**Figure 7.**
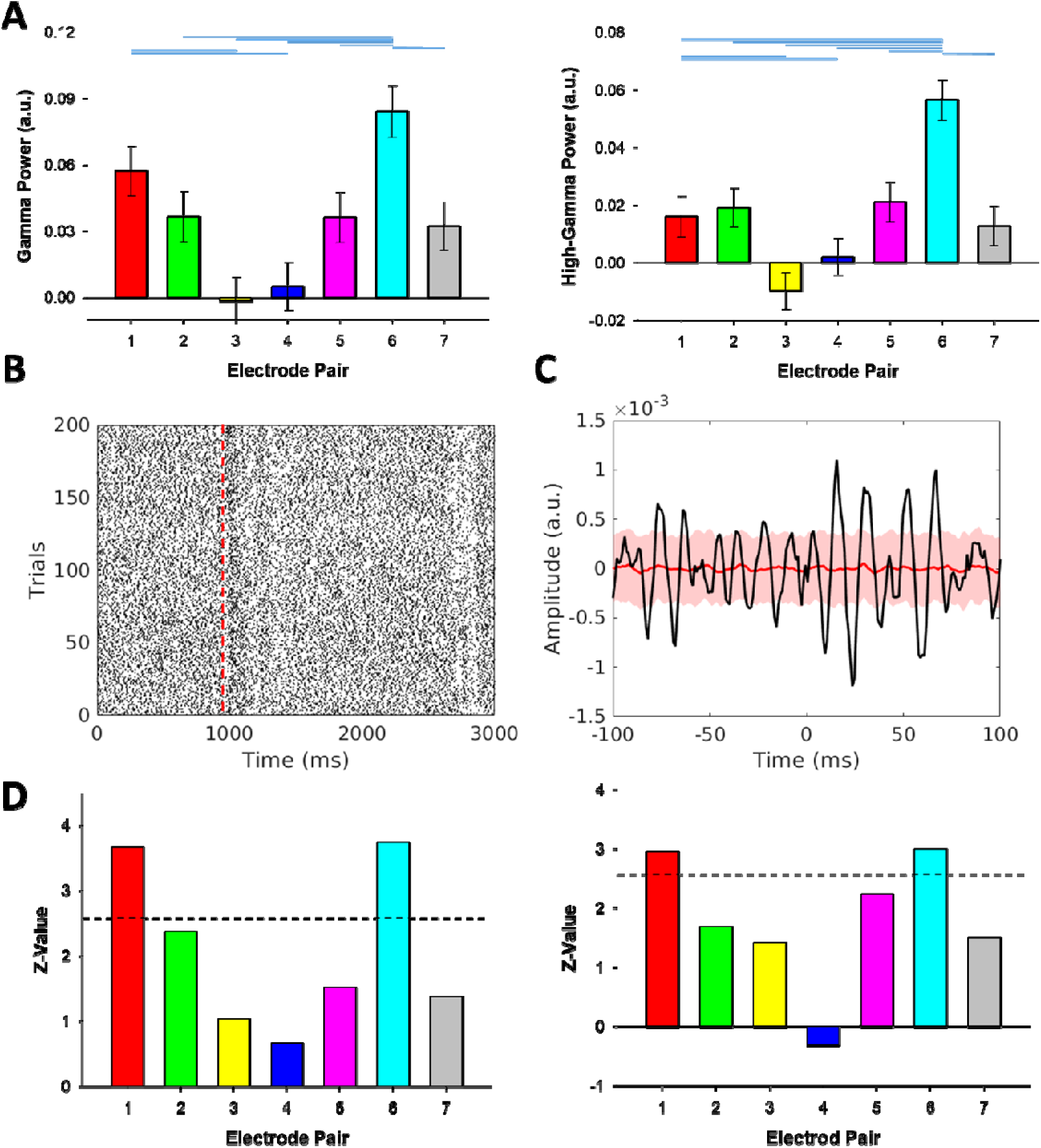
Layer-specific HG activities. (A, left) LHG power during pressure stimulation (0.5 to 1.5 s of stimulus onset). Electrode pairs 1, 2, 3, …, and 7 indicate bipolar-referenced electrode channels 1-2, 2-3, 3-4, …, and 7-8, respectively. Electrode pairs 1-3, 4-5, 6, and 7 were located in the layer VI, V, IV, and III, respectively. (A, right) HHG power during pressure stimulation. Horizontal blue lines above the bar graphs indicate the electrode pairs which show significant differences. Thin vertical lines indicate the standard error of the mean (s.e.m.). (B) Raster plot of neuronal spiking activities in the S1 during pressure stimulation. The red vertical line indicates stimulus onset. (C) Spike-triggered LFP (stLFP) during pressure stimulation (electrode pair 6). The red shaded area represents ±2 standard deviations (S.D.) of stLFP simulated by the Poisson process. The peak frequency was 66 Hz. (D) Normalized stLFP LHG (left) and HHG (right) band power at each electrode pair. Dashed horizontal lines indicate significance level (*Z* = 2.66, corrected).

## DISCUSSION

In this study, we showed that LHG and HHG activities have different functional roles in human S1. We suggest that HHG activity (70 to 150 Hz) represents the detail of surface geometry interacting with skin, while LHG (50 to 70 Hz) activity mainly represents the intensity of the sustained tactile stimulus. Additionally, we found that HHG activity to complex vibrotactile stimulus appears to be a mixture of HHG activities to flutter and vibration, indicating that there exists some degree of parallel processing between low and high mechanical frequencies in human S1. Finally, although we could not observe the power in the upper region of superficial layers, we found that both HHG and LHG power levels were highest in layer IV and that neuronal spiking activity in this layer was closely related to HG activity, suggesting that these high-frequency activities are potentially related to input information from the earlier stages of the somatosensory system, including the thalamus.

### LHG activity represents the intensity of sustained stimulation in S1

We showed that LHG activity increases with increasing the applied static force but HHG activity shows an inconsistent pattern. These results suggest that LHG, or narrowband gamma, activity has its own functional role distinct from HHG activity in S1. Previous vision studies revealed that narrowband gamma activity is related to light intensity (Saleem et al., 2017; Storchi et al., 2017). Similarly, we found that the intensity of sustained tactile stimulation modulates LHG activity. Considering the similarities between ECoG and LFP signals and the results of vision studies analyzing LFP (Dubey and Ray, 2020; Ray and Maunsell, 2011; Saleem et al., 2017), it is likely that LHG activities in S1 and V1 have similar processing mechanisms from a macroscopic perspective, even though the representing functions of the somatosensory and visual systems are completely different.

In this study, tHHG activity also increased with increasing applied force at the on/offset of stimuli. However, it is unlikely that this transient activity directly represents applied force during stimulation. Indeed, an ECoG study suggested that this activity is related to the time derivative of applied force (temporal dynamics of force) (Jiang et al., 2020). Similarly, although an MEG study using electrical stimulation suggested that gamma (30 to 100 Hz) amplitude encodes stimulus intensity in S1 (Rossiter et al., 2013), it appears to be due to the influence of the transient HG activity rather than narrowband gamma activity. Interestingly, the activation pattern of tHHG activity (on/offset-specific activation) is very similar to the neuronal firing pattern for the ramp-and-hold pressure stimulation (Callier et al., 2019).

### Neural activity to complex vibrotactile stimulation

In this study, we found that neural activity to complex vibrotactile stimuli was a mixture of neural activities to flutter and vibration. Although the convergence of submodality-specific inputs in S1 is known to exist (Pei et al., 2009; Saal and Bensmaia, 2014), our result implies that much of the initial neural processing for flutter and vibration occurs somewhat in parallel with each other, at least in S1. Assuming that the activity of S1 we measured mainly came from area 1, this is not surprising considering that nearly two-thirds of neurons in area 1 show afferent-specific neuronal activities (Pei et al., 2009). Conversely, this means that submodality convergence is not dominant in area 1 of S1. Perhaps such convergence occurs primarily in slightly higher-level somatosensory areas, including areas 2, 5 and the secondary somatosensory cortex (S2).

### HHG activity represents the dynamics of tactile stimulation

In our previous study, we observed that the sustained HG power during stimulation is different between coarse and fine textures (Ryun et al., 2017b). Additionally, a human ECoG study suggested that HG activity is related to the time derivative of applied force by observing the on/offset of grasping motion (Jiang et al., 2020). Here, our classification result indicates that the temporal dynamics of HHG activity contain tactile information about textured surfaces. In light of these findings, at least in tactile processing, HHG activity in S1 appears to be sensitive to subtle changes in skin deformation due to interaction between skin and objects. Further, although we distinguished between HHG and transient HG activities, both activities appear to be caused by the same mechanism, given the results that both activities were sensitive to the dynamics of mechanical stimuli.

### Origin of LHG and HHG activities

A vision study suggested that narrowband gamma oscillation is strongest in layer IV and reflects the signal flow from the thalamus to the cortex (Saleem et al., 2017). Furthermore, a recent study suggested that narrowband gamma oscillations (50 to 70 Hz) propagate and synchronize throughout the thalamocortical visual system in mice (Shin et al., 2023). Consistent with previous vision studies, in the current somatosensory study, we observed that LHG power was strongest in layer IV. We also found that neuronal spiking activity in this layer was closely related to LHG activity. In light of these findings, LHG activity (or narrowband gamma activity) may be induced by a thalamocortical circuit, such as the visual system, although we could not record the LHG power in the upper superficial layers. Meanwhile, HHG activity also showed the strongest power level in layer IV, indicating that HHG may also represent neuronal input activity. However, because 70-100 Hz gamma activity can be generated from superficial layers in S1 (although the sensation type is different) (Yue et al., 2020), further investigation is needed to confirm whether LHG and HHG activities actually occur in the same layer. Alternatively, it is also possible that HHG activities have multiple origins in S1.

### Limitations and further suggestions

In the current human study, we showed that texture types can be classified by temporal HG patterns in S1. However, we did not investigate the relationship between the physical properties of textured surfaces (e.g. grid size and vibration pattern of fingertip) and HG activities in S1. Previous primate studies showed that spatiotemporal firing patterns of neurons in S1 represent detailed physical properties of textured stimuli (Lieber and Bensmaia, 2019; Weber et al., 2013). Further studies are needed to reveal the relationship between HG activities and the physical properties of textured stimuli. In the current rat experiment, we were unable to record neuronal data from the upper part of the superficial layer. Additionally, although we determined cortical layers by estimating the insertion depth of the electrode bundle, we did not perform histological analysis to obtain additional information about the location of the electrode. Further studies using larger samples and histological analysis are required to determine the pattern of HG activity across the entire layer.

## MATERIALS AND METHODS

### Patient

Eight patients with drug-resistant epilepsy were included in this study. Patients underwent subdural ECoG implantation surgery before resection surgery. Pre-operative magnetic resonance (MR) and post-operative computed tomography (CT) images were obtained and used for MR-CT co-registration for localizing the electrodes. All experiments were approved by the Institutional Review Board (IRB) of Seoul National University Hospital (No.: 1610-133-803). All patients provided written informed consent before participation.

### Apparatus

A detailed description of the vibrotactile and texture stimulators is in our previous study (Ryun et al., 2023). For the complex vibrotactile stimulation, we designed a custom-made pin-point magnetic vibrator using a Mini- shaker (model 4810; Brüel and Kjær). A long plastic pin (2 mm in diameter; 100 mm in length) was mounted on the top of the Mini-shaker to stimulate an index fingertip. The plastic pin and Mini shaker were inserted in the stainless steel barrel to minimize electromagnetic and sound artifacts. For texture stimulation, a custom-made drum-type texture stimulator was used (**Figure S2**). The drum-type texture stimulator was designed to pseudo-randomly deliver eight different textures to the index finger. To measure the contact force on the skin, a small load cell was inserted under the plastic finger rest. The force data was simultaneously recorded using a DAQ device (USB 6002; National Instruments Corp., USA). A Lab Jack with micrometer precision positioned under the finger rest was used to control the contact force between the finger and the stimulator.

### Tasks

We delivered complex vibrotactile stimuli (flutter (32 Hz) + vibration (350 Hz)) to the index finger contralateral to the implantation site during ECoG recording. The maximum stimulus magnitude of the flutter and vibration were 100 μm and 50 μm, respectively to equalize the subjective intensity levels depending on the stimulus frequency (Mountcastle et al., 1990; Verrillo et al., 1969). The ratio between flutter and vibration was determined by these magnitudes (e.g. if flutter (75 %):vibration (25%), then flutter (90 μm) + vibration (15 μm)). The ratios used in this study were 99 (flutter):1 (vibration), 75:25, 50:50, 25:75, and 1:99. Throughout task sessions, all stimuli were passively delivered. Stimulus duration was 1.5 s, and the inter-trial interval was 2, 2.5, or 3 s. All stimuli were delivered pseudo-randomly. Paradigm was controlled by a custom-made software written in MATLAB (version 2018a, Mathworks, Natick, MA, USA).

For three patients, we pseudo-randomly and passively delivered eight texture stimuli to the patients’ index fingers. In this task, the contact force applied to the index finger has two conditions: light (20.09 ± 0.22 g wt, mean ± standard error) and heavy (93.42 ± 4.04 g wt) conditions. Texture materials were composed of coarse, intermediate and fine meshes, silk, film, sandpaper, and others (See **Figure S3**).

### Data Analysis

Data was recorded using Neuroscan or Neuvo (Neuroscan, Charlotte, NC, USA) with a 2000 Hz sampling rate. ECoG channels showing abnormal signals due to technical problems or epileptic activities were excluded from further analysis. A common average reference (CAR) was applied to the data for re-referencing. The data were notch-filtered with a zero-phase-lag infinite impulse response (IIR) filter to remove 60 Hz noise and its harmonics. We then performed epoching with a window of –2 to 3 s of stimulus onset. For time-frequency representation, a complex Morlet wavelet transform was applied to the single-trial data and then squared. The transformed single-trial data were normalized by the mean and standard deviation of the baseline (–1.5 to –0.5 s) signal at each frequency. To construct **Figure 1 (top)** and **3**, the transformed single-trial data were averaged across trials. To estimate similarity among various vibrotactile ratio conditions, we measured differences between 1:99 (flutter:vibration) and each stimulus condition or 99:1 and each stimulus condition using the following equation:

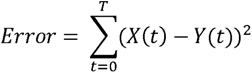

where *T* is the stimulus duration, *t* indicates the sample at each time point, and *X* and *Y* are the HG power levels of two different conditions. The error values were then normalized by the value of the maximum error condition. To explore that HHG activity for complex vibrotactile stimuli can be expressed as a superposition of HG activity for flutter and vibration-only conditions, we first averaged HHG time series of each condition across all subjects and then found the values that minimize the error of the following equation:

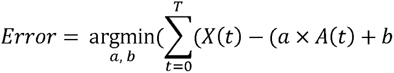

where *T* is the stimulus duration, *t* indicates the sample at each time point, *X* is the HHG power level of complex vibrotactile stimulation (e.g., 75:25, 50:50, and 25:75), *A* and *B* are the HHG power levels of flutter and vibration-only conditions, respectively, and the values *a* and *b* indicate the coefficients of *A* and *B*, respectively. To calculate the power of each HG component (higher component (70 to 150 Hz and 250 ms to 1.5 s), lower component (50 to 70 Hz and 250 ms to 1.5 s), and transient HG (70 to 150 Hz and 0 to 250 ms)), the power values over the corresponding frequency and time ranges were averaged. To test the significance among each condition, we performed an independent two-sample t-test. For multiple comparison corrections, Bonferroni correction was performed.

For texture classification, we first extracted features from the transformed single-trial data. We averaged the power in the range of 50-70 Hz and 70-150 Hz for LHG and HHG respectively. We then divided the HG power time series into five bins (250 ms each), and the average power levels of each bin were used for the feature. For transient HG (70 to 150 Hz at 0-250 ms) condition, we also divided the power time series into 5 bins (50 ms each).

For classification, we designed the multi-layer perceptron (MLP) classifier by using the Keras library with Tensorflow 2.5 on an NVIDIA GeForce RTX 4070Ti GPU. We sequentially construct an MLP architecture consisting of an input layer, a fully connected layer of 10 nodes, a 50 % dropout layer, and a dense layer of eight outputs. We used the ‘categorical cross entropy’ option for the loss function and used the ‘ADAM’ optimizer. The number of epochs was 400, and the batch size was 32. We used 8-fold cross-validation to evaluate classification performance. The whole process was repeated five times, and thus we evaluated the classification accuracy using the test dataset 40 times.

### Rat Experiment

In the rat experiment, we inserted a microelectrode bundle (silicon probe from Plexon Inc., USA) with eight vertical channels into the hindlimb area of the rat’s left somatosensory cortex (electrode diameter of 20 μm; inter-electrode distance of 200 μm). Electrodes 1-3, 4-5, and 6-7 were located in the layers VI, V, and IV, respectively. Electrode 8 was located in the lower part of the superficial layer (approximately layer III). We estimated the layer corresponding to each electrode based on the insertion depth of the electrode bundle (Defelipe, 2011). We delivered mechanical pressure stimuli to the right hindlimb of an anesthetized (with isoflurane) rat by using the wood pin attached to the servo motor. The Servo motor was controlled by the Arduino microcontroller unit (MCU). The stimulus duration was 2 s, and we repeated the mechanical stimulation 200 times with an inter-stimulus interval of 3 s in one session (6 sessions total). For comparison, a similar experiment was performed on humans. We used a Von Frey filament to manually deliver static pressure to the index finger. Contact force, stimulus duration, and the number of trials were approximately 300 g wt., 1 s, and 40 trials, respectively.

Neuronal data were recorded by the Plexon acquisition system (Plexon Inc., USA) at a 20k Hz sampling rate. For spike detection and sorting, we used WaveClus toolbox (version 2.5). The spike detection threshold was 5.5 standard deviations of the magnitude of the background noise signal. For unsupervised spike clustering, we used the superparamagnetic clustering method (n = 14 neurons from 8 electrodes) (Quiroga et al., 2004). To analyze local field potentials, the raw data were re-referenced using the bipolar reference method (e.g., 1-2, 2-3, …, 7-8). The data were low-pass filtered at 200 Hz and then downsampled to 1000 Hz. We used a complex Morlet wavelet transform to calculate time-frequency representation. To calculate the power levels of HHG and LHG activities, we averaged the power level data across the frequency (50 to 70 Hz and 70 to 150 Hz for LHG and HHG power, respectively) and time 0.5 s to 1.5 s of stimulus onset). For significance testing, we used one-way ANOVA and then performed the Tukey-Kramer test for post-hoc analysis. To calculate spike-triggered local field potential (stLFP), LFPs for each electrode pair were aligned by the onsets of neuronal spikes (window size: ±500 ms) during pressure stimulation and then averaged across spike events. For visualization (Figure 6E), stLFPs were high-pass filtered at 40 Hz. To test the significance of oscillatory activities in stLFPs, we created spike trains during stimulation by utilizing the Poisson process with the same firing rate as the actual ones. Then we calculated stLFPs for the created spike events and repeated this procedure 200 times to obtain the LHG and HHG band power distributions of the stLFPs. We used the “normfit” Matlab function to extract the Z-values of the real oscillatory activity. We performed this analysis for all possible neuron-LFP pairs within each electrode pair. Significant neuron-LFP pairs were only observed for electrode pairs 1 and 7 among all possible pairs.

## ACKNOWLEDGMENTS

This research was supported by the Alchemist Brain to X (B2X) Project funded by Ministry of Trade, Industry and Energy (20012355, NTIS: 1415181023).

## AUTHOR CONTRIBUTIONS

S. Ryun and C. K. Chung designed and conducted the study. C. K. Chung performed the surgeries. S. Ryun designed the tactile devices and experimental paradigm. S. Ryun carried out experiments and performed data analysis. S. Ryun and C. K. Chung wrote the paper.

