## Supplementary Information for "Distinct Functional Roles of Narrow and Broadband High-Gamma Activities in Human Primary Somatosensory Cortex"

**Table 1.** Demographics of the patients

| Patient No. | Age | Sex | Number of Electrodes | Task | Electrodes location | Diagnosis |
| --- | --- | --- | --- | --- | --- | --- |
| 1 | 30 | F | 96 | Texture | Right | TLE and PLE |
| 2 | 40 | F | 48 | Vibrotactile | Right | FLE |
| 3 | 40 | M | 62 | Vibrotactile, Texture | Left | Hemispheric epilepsy |
| 4 | 23 | F | 48 | Vibrotactile | Left | TLE |
| 5 | 24 | M | 62 | Vibrotactile, Texture | Left | FLE |
| 6 | 21 | M | 24 | Vibrotactile | Right | TLE |
| 7 | 29 | F | 54 | Vibrotactile | Right | TLE |
| 8 | 58 | M | 52 | Vibrotactile | Left | TLE |

Abbreviation: M = male, F = female, TLE = temporal lobe epilepsy, PLE = parietal lobe epilepsy, FLE = frontal lobe epilepsy.

**Supplementary Figures**

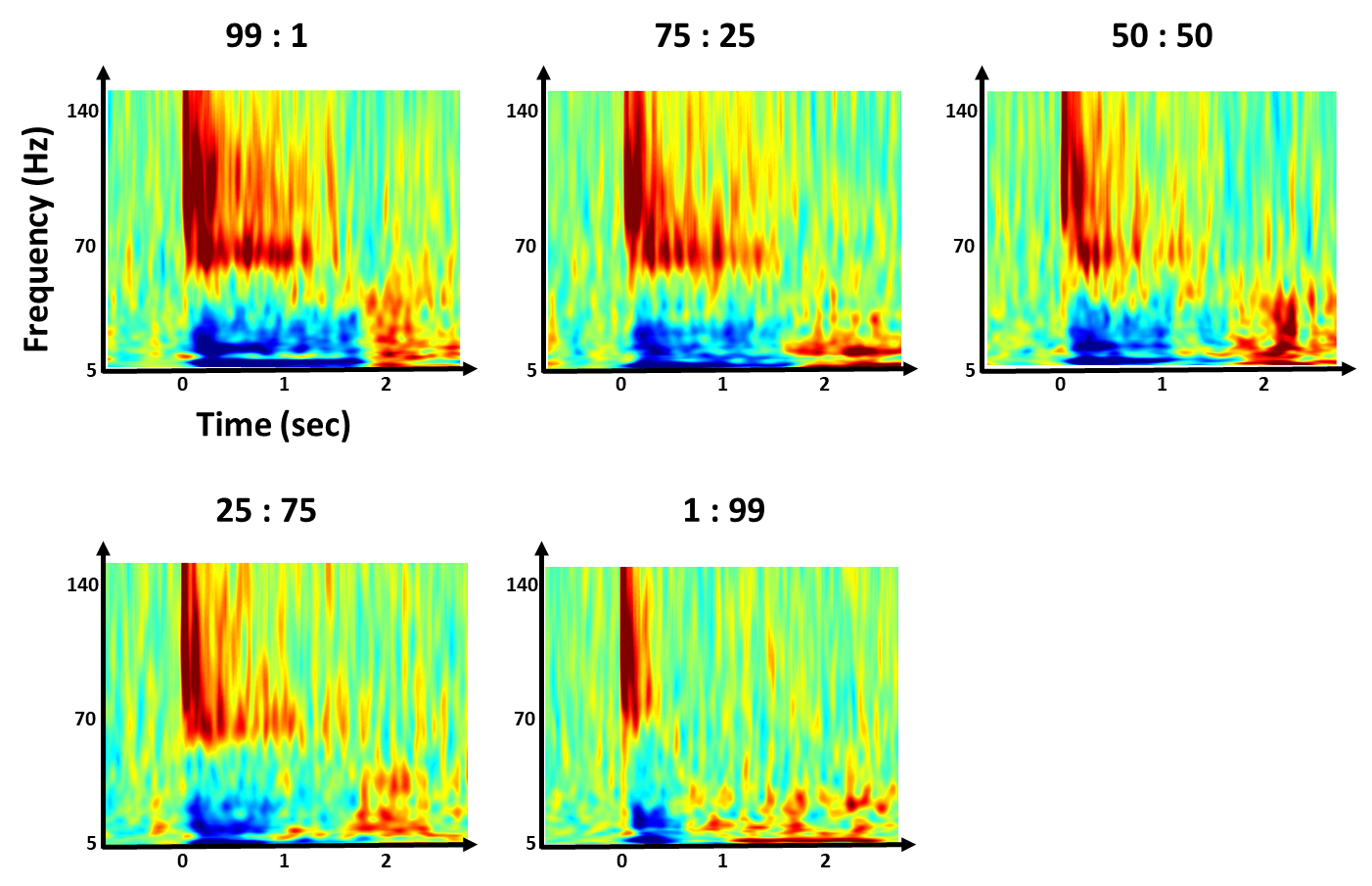

**Figure S1.** Time-frequency maps under various flutter (32 Hz) + vibration (350 Hz) conditions from subject 3

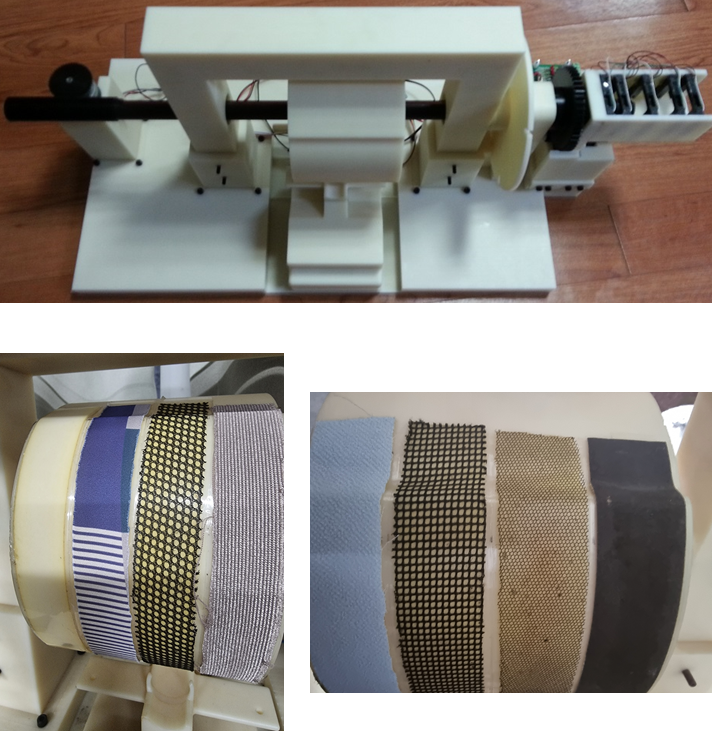

**Figure S2.** Drum-type texture stimulator.

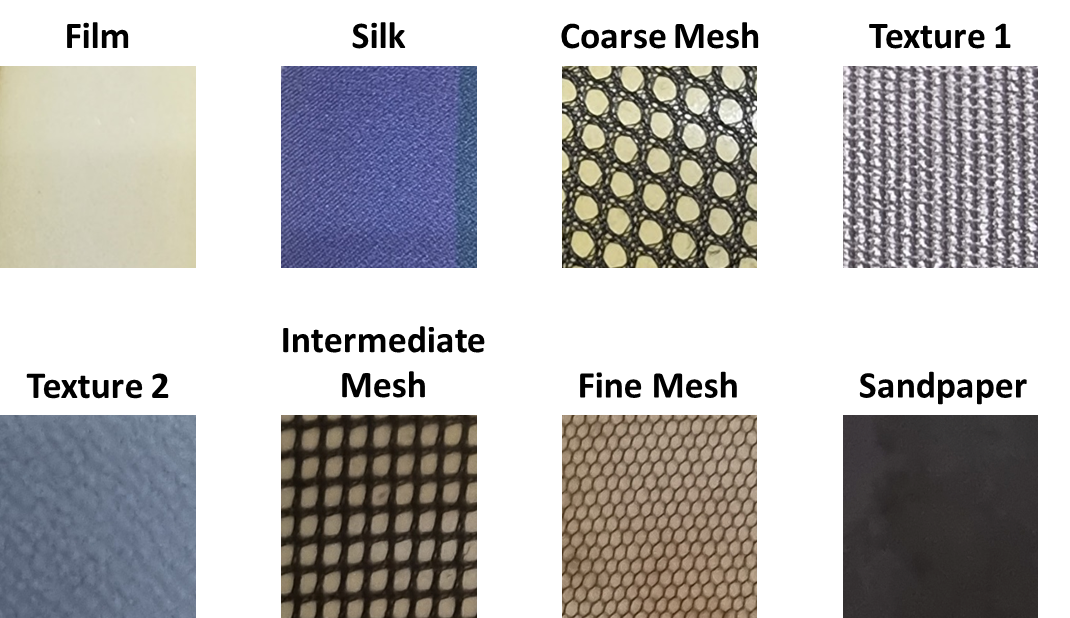

**Figure S3.** Eight textures used in this study.
